# EEG signals index a global signature of arousal embedded in neuronal population recordings

**DOI:** 10.1101/2020.08.03.235283

**Authors:** Richard Johnston, Adam C. Snyder, Rachel S. Schibler, Matthew A. Smith

**Affiliations:** Department of Biomedical Engineering, Carnegie Mellon University, Pittsburgh, USA; Neuroscience Institute, Carnegie Mellon University, Pittsburgh, USA; Center for the Neural Basis of Cognition, Carnegie Mellon University, Pittsburgh, USA; Department of Brain and Cognitive Sciences, University of Rochester, Rochester, USA; Department of Neuroscience, University of Rochester, Rochester, USA; Center for Visual Science, University of Rochester, Rochester, USA; Harvey Mudd College, California, USA

## Abstract

Electroencephalography (EEG) has long been used to index brain states, from early studies describing activity during visual stimulation to modern work employing complex perceptual tasks. These studies shed light on brain-wide signals but lacked explanatory power at the single neuron level. Similarly, single neuron studies can suffer from inability to measure brain-wide signals. Here, we combined these techniques while monkeys performed a change detection task and discovered a link between EEG and a signal embedded in spiking responses. This ‘slow drift’ was associated with arousal: decreases in pre-stimulus α power/increases in P1 amplitude were accompanied by :1) increases in pupil size, false alarm rate and saccade velocity; and 2) decreases in microsaccade rate and reaction time. These results show that brain-wide EEG signals can be used to index modes of activity acquired from direct neural recordings, that in turn reflect global changes in brain state that influence perception and behavior.

## Introduction

For decades, researchers have investigated how the spiking responses of single cortical neurons relate to performance on decision-making [1], attention [2] and working memory [3] tasks. Interactions between pairs of neurons have also been studied extensively since technological advances in neural recording systems (e.g. microelectrode arrays and two-photon imaging) made it possible to monitor the activity of neural populations simultaneously [4–6]. At the same time, it is becoming increasingly apparent that major insight about the neurobiological basis of cognition can be gained if one goes beyond studying single and pairwise statistics [7–14]. Furthermore, it has been suggested that low-dimensional neural activity patterns can be used to index global brain states, which influence performance on cognitive tasks. For example, Stringer et al. [15] applied principal component analysis (PCA) to data recorded from more than 10,000 neurons in the mouse and found that fluctuations in the first principal component were associated with a host of arousal-related variables including whisking, pupil size, and running speed. Musall et al. [16] found that uninstructed movements, which themselves may occur at varying frequency based on arousal, were related to brain-wide activity in the mouse. In our own work in macaque monkeys, we have reported a brain wide ‘slow drift’ of neural activity [17] which is correlated with a distinctive pattern of eye metrics that is strongly indicative of changes in arousal [18]. However, it is unknown if they are associated with other arousal-related variables that can be measured in a rapid, accurate and non-invasive manner.

Spontaneous (i.e., pre-stimulus) oscillations in the α frequency band (∼8 – 12Hz) are associated with global changes in brain state. For example, research in humans using EEG and magnetoencephalography (MEG) has shown that the likelihood of detecting a near-threshold visual stimulus increases when pre-stimulus oscillations in the α band decrease [19–26]. The principles of signal detection theory [27] dictate that changes in behavior on detection tasks can arise due to changes in sensitivity or response criterion [28–30]. To that end, recent work has shown that a decrease in pre-stimulus α power is associated with increased hit rate and false alarm rate [31,32]. These results point to link between pre-stimulus α power and response criterion, a key component of signal detection theory, which is modulated, at least in part, by subcortical structures that control arousal levels [31,33]. In addition, pre-stimulus α power is known to be correlated with eye metrics including pupil size [34,35] and reaction time [35–38] such that decreased pre-stimulus α power is accompanied by increased pupil size and decreased reaction time. These findings raise the possibility that other EEG signals such as the P1 component of the visually evoked potential (VEP) might also be associated with arousal.

Several studies have established a link between early components of the VEP and changes in global brain state. In contrast to pre-stimulus α oscillations, P1 amplitude is significantly larger on trials in which a weak visual stimulus is detected in comparison to when it is not detected [19,25,39,40]. Furthermore, studies that have explored the effects of spatial attention on early components of the VEP have shown that P1 amplitude is negatively correlated with reaction time [41]. In light of these results, and the well-known involvement of oscillatory brain activity in stimulus-evoked responses [42], one might expect a negative relationship between pre-stimulus α oscillations and early VEP components. There is some evidence to suggest that this is the case. For example, Iemi et al. [43] found that the amplitude of the C1 and N150 increased when pre-stimulus power in the α frequency band decreased. These results suggest that pre-stimulus α power and early components of the VEP (e.g. P1 amplitude) can be used to index global changes in arousal. However, it is currently unclear how to link these EEG signals to underlying patterns of neural activity in the cortex.

In this study, we explored if pre-stimulus oscillations in the α frequency band and P1 amplitude are associated with slow drift in visual cortex. We simultaneously recorded EEG from the scalp and spiking activity from populations of neurons in V4 of two monkeys while they performed an orientation-change detection task (Figure 1A). Results showed that neural slow drift was associated with a pattern that is indicative of changes in the subjects’ arousal levels over time. For example, decreases in pre-stimulus power specifically in the α band (i.e., not other frequencies) and increases in P1 amplitude were accompanied by 1) increases in pupil size, false alarm rate and saccade velocity; and 2) decreases in microsaccade rate and reaction time. These findings are important because they reveal that spontaneous/evoked components of the EEG signal recorded non-invasively on the scalp index low-dimensional patterns of neural activity acquired from microelectrode array recordings in the brain. They support previous research [17,18] showing that slow drift is associated with gradual changes in arousal over time, and provide a strong link between global measurements made across recording modalities and species.

**Figure 1.**
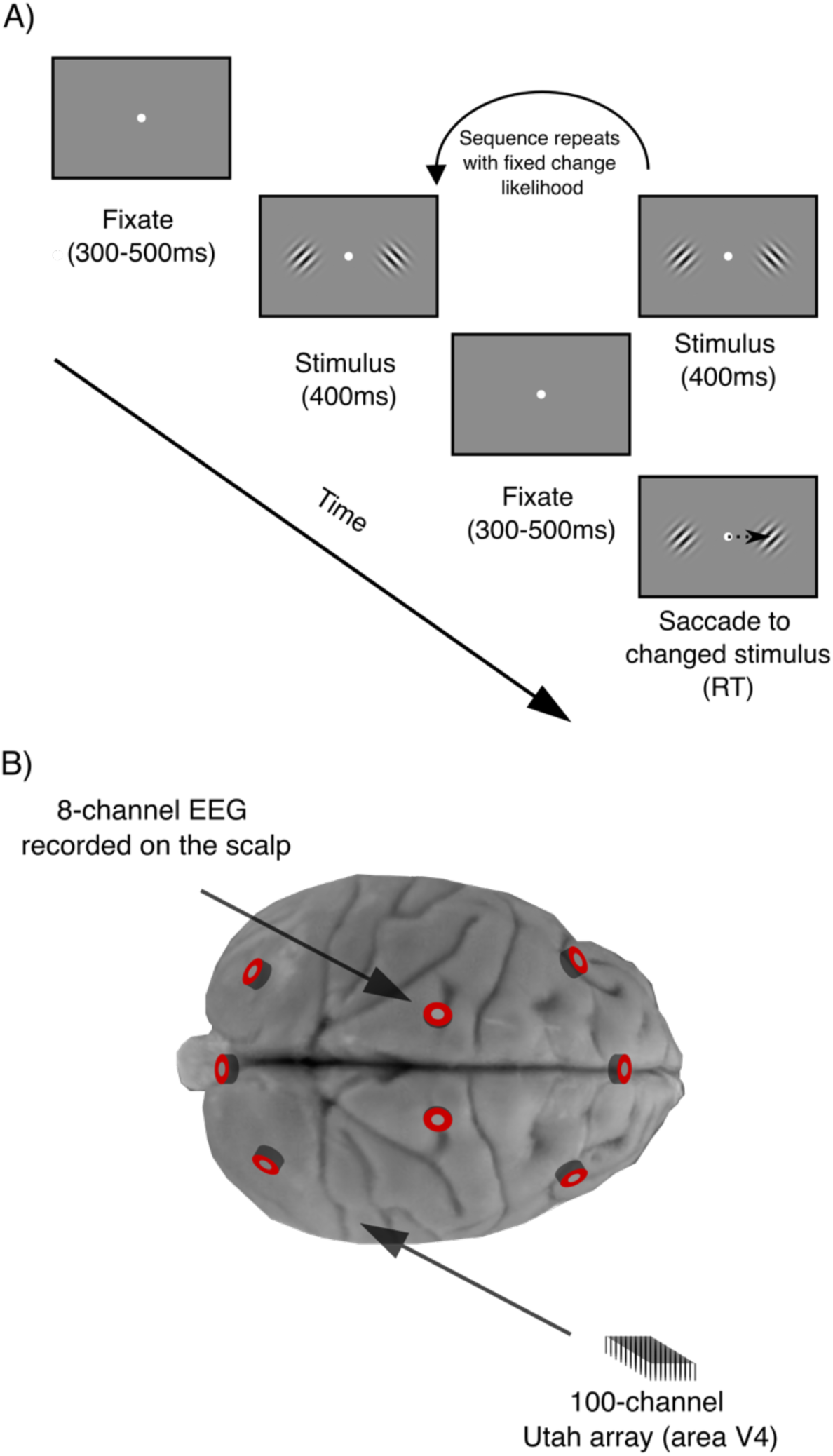
Experimental methods. (A) Orientation-change detection task. After an initial fixation period, a sequence of stimuli (orientated Gabor pairs separated by fixation periods) was presented. The subjects’ task was to detect an orientation change in one of the stimuli and make a saccade to the changed stimulus. (B) Electrophysiological recordings. We simultaneously recorded: 1) spiking responses of populations of neurons in V4 using 100-channel microelectrode (‘Utah’) arrays; and 2) EEG from 8 electrodes positioned on the scalp.

## Results

To determine if spontaneous and evoked components of the EEG signal can provide insight into the internal brain state associated with slow drift, we trained two macaque monkeys to perform an orientation change-detection task in which pairs of stimuli were repeatedly presented (Figure 1A). Spiking responses of populations of neurons in visual cortex (V4) were recorded using 100-channel “Utah” arrays as well as EEG on the scalp (Figure 1B). On each trial, we calculated: 1) the mean amplitude of oscillations in different frequency bands during each pre-stimulus period (300-500ms) using a fast Fourier transform (FFT); and 2) the VEP during each 400ms stimulus period (see Methods). Previous research has shown that pre-stimulus α power and stimulus-evoked P1 amplitude across a wide range of frontal, midline and occipital electrode sites are associated with improved performance on detection tasks [19,22,31]. Hence, the signals from all 8 EEG electrodes (referenced online to the head post, see Methods) were averaged prior to computing the FFTs and VEPs. We focused primarily on pre-stimulus power in the α frequency band and P1 amplitude as they capture spontaneous (i.e., pre-stimulus) and evoked aspects of the EEG signal and have been linked to global changes in brain state. However, we also measured pre-stimulus beta (β) and gamma (γ) oscillations to exclude the possibility that slow drift was associated with aperiodic (i.e. broadband) changes in spectral power over time (1/f noise) [43–46].

### Correlation between pre-stimulus α power and P1 amplitude

First, we explored the relationship between pre-stimulus α power and the amplitude of the P1 component of the VEP. As described above, these two signals appear to operate in an antagonisticmanner on detection tasks with decreases in pre-stimulus α power/increases in P1 amplitude being associated with improved performance [19–26,39,40]. Hence, we hypothesized that these two large-scale EEG signals would be negatively correlated over time. To investigate the relationship between pre-stimulus α power and P1 amplitude, EEG data for each session was binned using a 30-minute sliding window stepped every 6 minutes (Figure 2A and Figure 2B). The width of the window, and the step size, were chosen to isolate slow changes over time based on previous studies we performed [17,18]. An example session is shown in Figure 2C. In support of our hypothesis, pre-stimulus power in the α frequency band was negatively associated with P1 amplitude.

**Figure 2.**
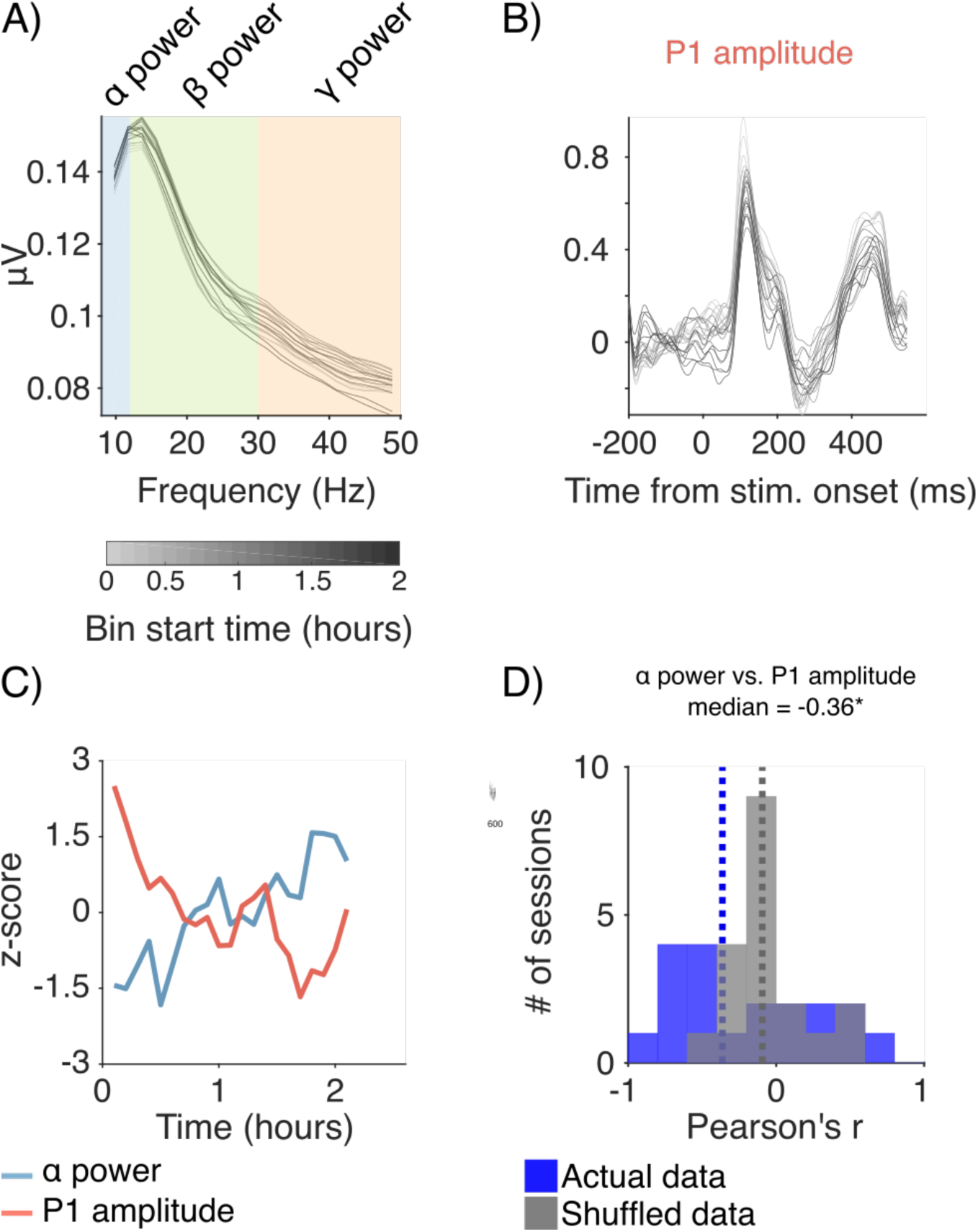
Isolating slow fluctuations in spontaneous and evoked EEG signals. (A) Spontaneous EEG signals. On each trial, we calculated the mean amplitude of oscillations in distinct frequency bands for each pre-stimulus period using a FFT. The data was then binned using a 30-minute sliding window stepped every 6 minutes. Each bin is represented by a shaded grey line. Note that the duration of the fixation period was held constant at 300ms in this plot for illustration purposes only. In our analysis, we used the full duration of the fixation period, which was randomized between 300 to 500ms. (B) Evoked EEG signals. On each trial, we calculated a VEP for each stimulus presentation. The data was then binned using the same 30-minute sliding window so that direct comparisons could be made between spontaneous and evoked EEG signals. Each bin is represented by a shaded grey line. (C) Example session from Monkey 1 showing that pre-stimulus α power was negatively correlated with P1 amplitude over time. Each metric has been z-scored for illustration purposes. (D) Histogram showing distributions of Pearson’s r values (black) across sessions between pre-stimulus α power and P1 amplitude. Actual distributions were compared to shuffled distributions (grey) using two-sided permutation tests (difference of medians). Median r values are indicated by dashed lines. p < 0.05*, p < 0.01**, p < 0.001***. See Figure S1 for additional example sessions from both monkeys.

Next, we explored if a similar pattern was present across sessions. We computed correlations (Pearson product-moment correlation coefficient) between pre-stimulus power in the α band and the amplitude of the P1 component for each session and compared the distribution of Pearson’s r values obtained to shuffled distributions using permutation tests (two-sided, difference of medians). Consistent with the pattern of results observed in several individual sessions for Monkey 1 and Monkey 2 (Figure S1), we found that pre-stimulus α power was significantly and negatively correlated with P1 amplitude (Figure 2D, median r = −0.36, p = 0.024). These results motivated us to ask if pre-stimulus α power and P1 amplitude are correlated with other task performance and eye metrics that are often taken to be hallmarks of arousal e.g. hit rate, false alarm rate, pupil size, microsaccade rate, reaction time and saccade velocity. A direct prediction of our results is that an opposite pattern should emerge when these measures are correlated with pre-stimulus α power and P1 amplitude.

### Correlating pre-stimulus α power/P1 amplitude with other arousal-related variables

In order to directly compare pre-stimulus α power and P1 amplitude with a host of arousal-related variables we binned the task performance (hit rate and false rate) and eye metrics (pupil size, microsaccade rate, reaction time, saccade velocity) in the manner described above using a 30-minute sliding window stepped every six minutes. We computed Pearson product-moment correlations between these metrics for each session and we compared the subsequent distributions of Pearson’s r values to shuffled distributions using permutation tests (two-sided, difference of medians). First, we found that the amplitude of pre-stimulus oscillations in the α band was negatively correlated with false alarm rate (Figure 3A, median r = −0.39, p < 0.001). No significant correlation was found between pre-stimulus α power and hit rate (median r = 0.04; p = 0.783) but false alarm rate was itself positively correlated with hit rate across sessions (Figure S2, median r = 0.49, p < 0.001). Hence, our results support an emerging body of work showing that the amplitude of pre-stimulus α oscillations is associated with changes in response criterion as opposed to sensitivity [32,43]. In addition, a negative correlation was found across sessions between pre-stimulus α power and pupil size in Monkey 1 (median r = −0.32, p = 0.037), but no such correlation was found in Monkey 2 (median r =0.07 p = 0.623). We found a significant (p<0.05) correlation between pre-stimulus α power and pupil size in 7/20 sessions for Monkey 1 and 2/16 sessions for Monkey 2. Across sessions and monkeys, we found positive correlations of pre-stimulus α power with reaction time (Figure 3A, median r = 0.26, p = 0.012), and P1 amplitude with saccade velocity (Figure 3B, median r = 0.21, p = 0.026). Negative correlations of pre-stimulus α power with saccade velocity (median r = −0.27, p = 0.002), and P1 amplitude with microsaccade rate (median r = −0.38, p = 0.008) were also observed. No significant correlation was found between P1 amplitude and reaction time (median r = 0.08, p = 0.728), but reaction time was itself negatively correlated with saccade velocity across sessions (Figure S2, median r = −0.52, p < 0.001). Taken together, these results demonstrate that slow time scale changes in spontaneous and evoked EEG signals were accompanied by changes in the subjects’ arousal levels over time. In particular, decreases in pre-stimulus α power and increases in P1 amplitude were accompanied by increases in false alarm rate and saccade velocity as well as decreases in microsaccade rate and reaction time.

**Figure 3.**
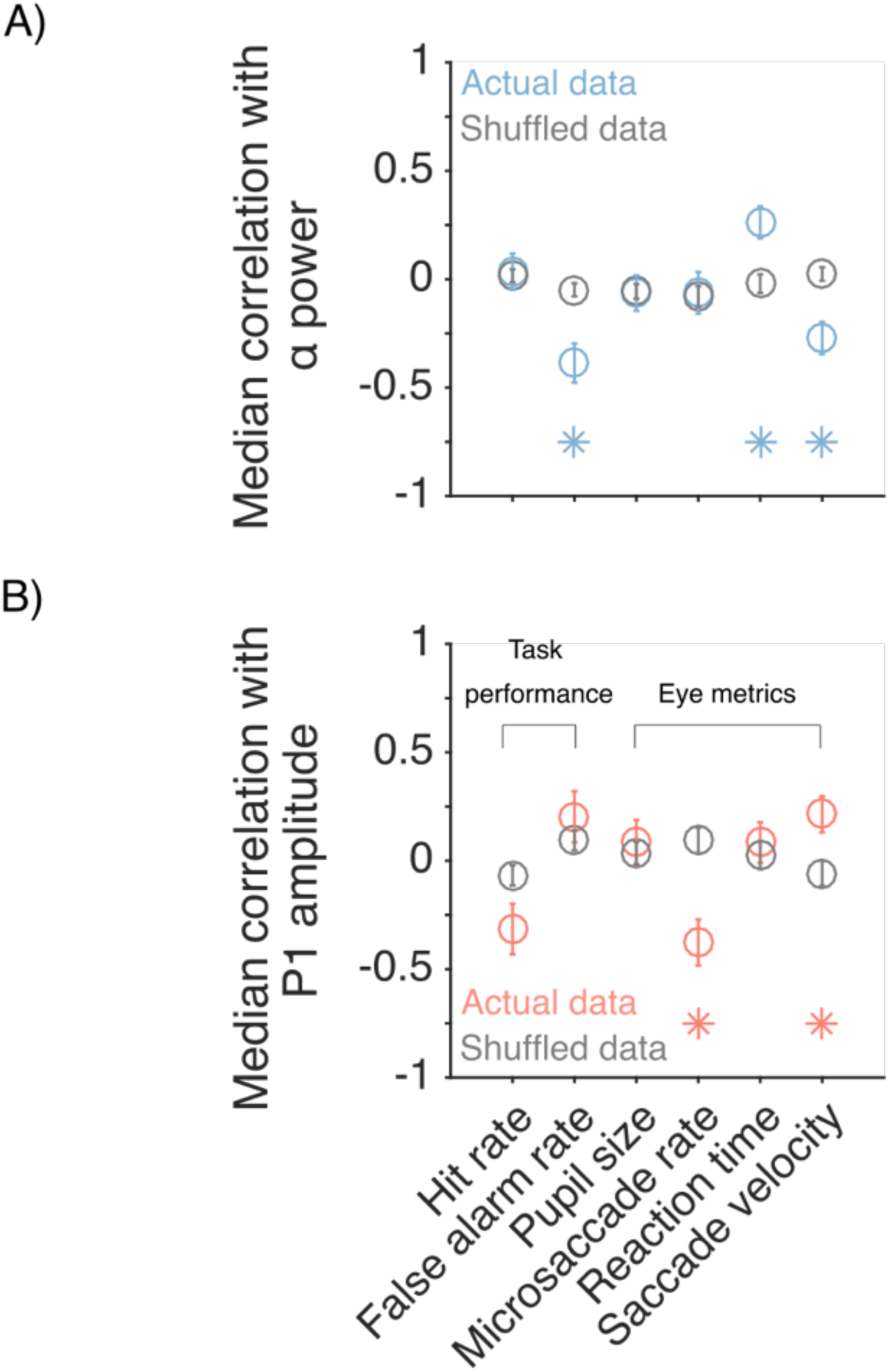
Correlations between EEG signals and other arousal-related variables. (A) Pre-stimulus α power. Each blue point corresponds to the median r value across sessions between pre-stimulus α power and a given metric. In contrast, grey pints represent median r values computed using shuffled data. Actual distributions of r values were compared to shuffled distributions using two-sided permutation tests (difference of medians). Asterisks indicative a significant effect with a p value at least < 0.05. See Figure S2 for histograms showing distributions of r values (black) across sessions between: A) hit rate and false alarm rate; and B) reaction time and saccade velocity.

### Correlations between pre-stimulus α power, P1 amplitude and slow drift

As described above, we and others have shown that pre-stimulus α power and P1 amplitude are related to global changes in arousal. Therefore, we were keen to explore if these indirect and non-invasive measures of neural activity are associated with low-dimensional modes of neural activity acquired directly from the spiking activity of a population of neurons. Previously, we reported a co-fluctuation of neural activity in macaque visual and prefrontal cortex [17]. This ‘slow drift’ was related to the subject’s tendency to make impulsive decisions in a change detection task, ignoring sensory evidence (false alarms), and multiple eye metrics that together are indicative of global changes in brain state [18]. Here we sought to determine if pre-stimulus α power and P1 amplitude are associated with slow drift.

To calculate slow drift, we binned spike counts in V4 using the same 30-minute sliding window described above and used in previous research (Figure 4A, see Methods). We then applied principal component analysis (PCA) to the data and estimated slow drift by projecting the binned residual spike counts along the first principal component (i.e. the loading vector that explained the most variance in the data). Because the sign of the loadings in PCA is arbitrary [47], the correlation between slow drift and a given variable can be positive or negative. This would not affect the overall pattern of results if one were solely interested in total variance explained. However, we were interested in whether slow drift was associated with a characteristic pattern of EEG activity that we and others have shown to be indicative of changes in the subjects’ arousal levels over time i.e. decreased pre-stimulus α power and increased P1 amplitude (Figure 2 and Figure 3). Hence, the sign of the correlation between slow drift and the spontaneous/evoked EEG signals was critical. In order to establish a common frame of reference across sessions, and preserve the sign of the correlations, we constrained the slow drift for each session to have the same relationship to the spontaneous and evoked activity of the neurons. That is, the slow drift was flipped if the mean projection value for evoked responses recorded during stimulus periods was less than that for spontaneous responses recorded during pre-stimulus periods i.e., if the relationship that was naturally observed for unprojected data did not hold true [18]. This served to align the data across sessions such that relatively large slow drift values were associated with higher firing rates. To put it another way, a negative correlation between pre-stimulus α power and slow drift would be indicative of a pattern in which decreased pre-stimulus α power would lead to higher firing rates. Our hypothesis predicts that the opposite should be true for P1 amplitude.

**Figure 4.**
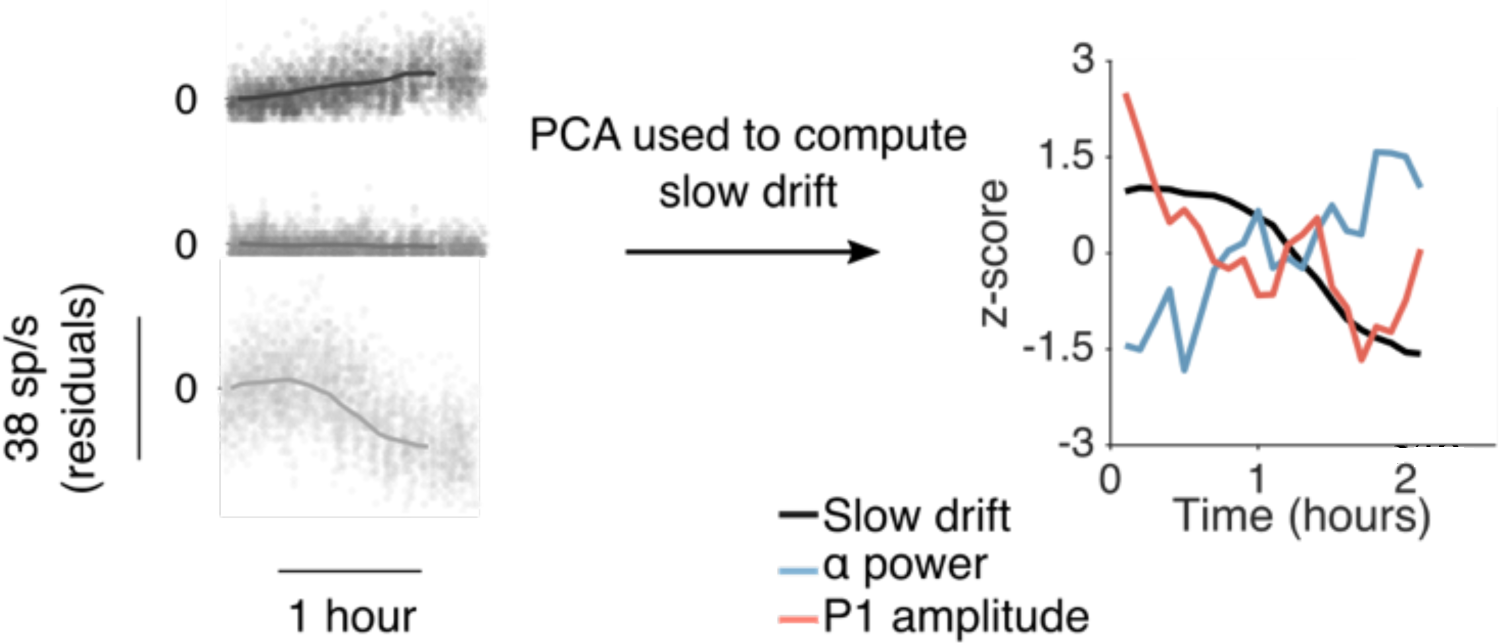
Calculating slow drift. (A) Three example neurons from a single session (Monkey 1). Each point represents the mean residual spike count during a 400ms stimulus period. The data was then binned using a 30-minute sliding window stepped every six minutes (solid line) so that direct comparisons could be made with the EEG signals. PCA was used to reduce the dimensionality of the data and slow drift was calculated by projecting binned residual spike counts along the first principal component. (B) Example session from Monkey 1 showing that slow drift was negatively correlated with pre-stimulus α power and positively correlated with P1 amplitude. Each metric has been z-scored for illustration purposes.

**Figure 5.**
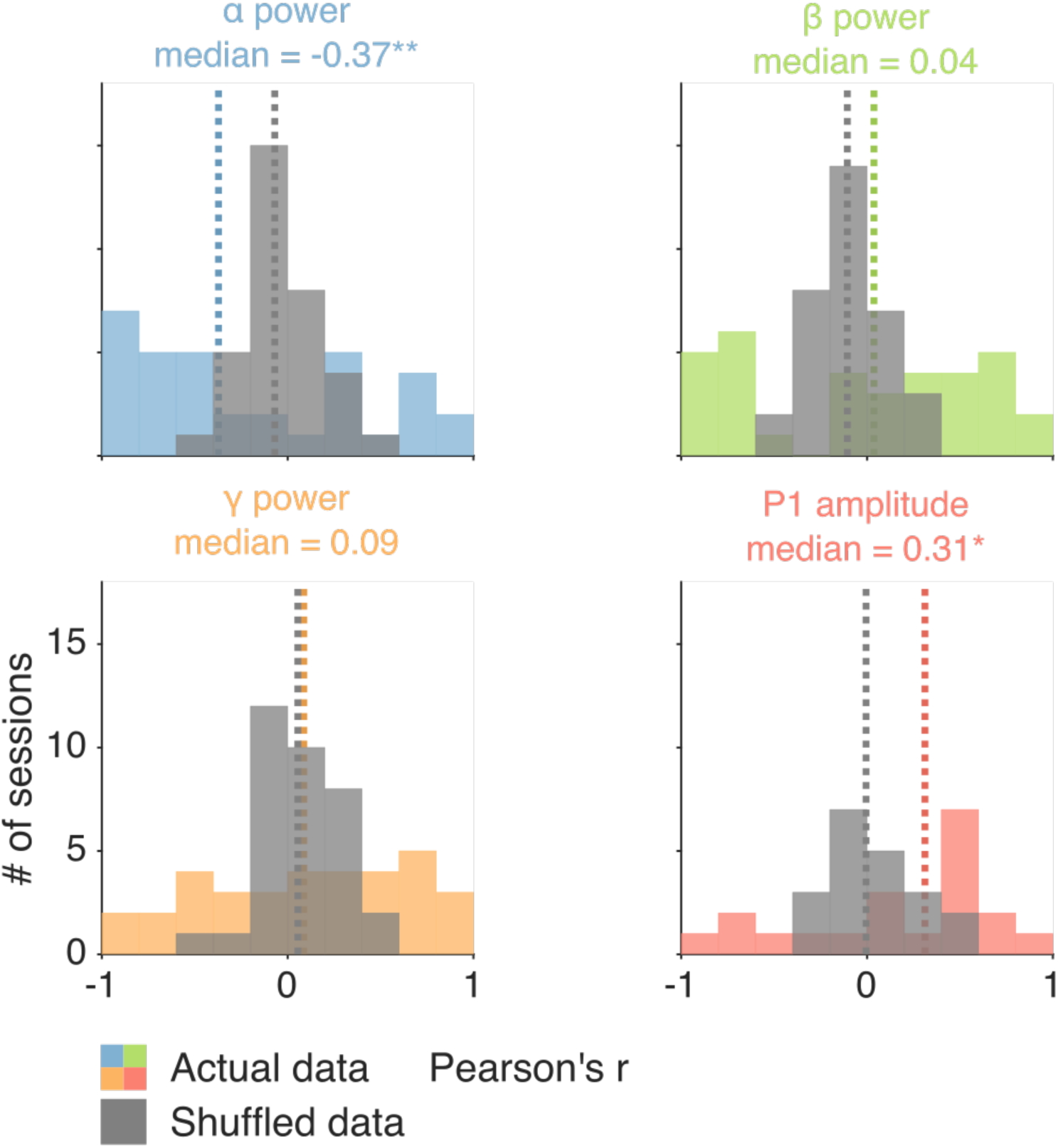
Correlations between EEG signals and slow drift in visual cortex. Histogram showing distributions of r values (colors) across sessions between slow drift and spontaneous/evoked EEG signals. Actual distributions were compared to shuffled distributions (grey) using two-sided permutation tests (difference of medians). Median r values are indicated by dashed lines. p < 0.05*, p < 0.01**, p < 0.001***. See Figure S3 for histograms showing distributions of r values across sessions between slow drift and aperiodic-adjusted power in different frequency bands.

We computed the slow drift of the neuronal population in each session using the above-mentioned method, and then compared it to our two spontaneous and evoked EEG measures: pre-stimulus α power and P1 amplitude. An example session is shown in Figure 4B (same sessions as in Figure 2C). In support of our hypothesis, a characteristic pattern was found in which slow drift was negatively associated with pre-stimulus α power and positively associated with P1 amplitude. Next, we investigated if a similar pattern was found across sessions.

We computed the Pearson product-moment correlation coefficient between pre-stimulus α power/P1 amplitude and slow drift for each session. We then compared the actual distribution of Pearson’s r values to shuffled distributions using permutation tests (two-sided, difference of medians). Because the slow drift was aligned across sessions based on the neural activity alone and not the EEG metrics, the shuffled distributions were centered on a correlation value of zero. We also computed correlations between slow drift and pre-stimulus oscillations in the beta (β) and gamma (γ) frequency bands to rule out the possibility that slow drift was associated with aperiodic (i.e. broadband) changes in spectral power over time (1/f noise) [43–46]. If this was the case, one might expect a significant correlation between slow drift and the amplitude of pre-stimulus oscillations in the α, β and γ frequency bands.

Consistent with the pattern of results observed in several individual sessions for Monkey 1 and Monkey 2 (Figure S1), we found that slow drift was negatively correlated with pre-stimulus α power (median r = −0.38, p = 0.004), whereas it was positively correlated with P1 amplitude (median r = 0.31, p = 0.027). No significant correlation was found between slow drift and the amplitude of pre-stimulus power in the β (median r = 0.04, p = 0.078) or γ (median r = 0.09, p = 0.621) bands. Although these findings suggest that slow drift was not associated with changes in 1/f noise over time [43–46] we carried out an additional analysis in which aperiodic signals were removed prior to computing mean oscillatory power in different frequency bands. Importantly, the overall pattern of results remained the same when slow drift was correlated with aperiodic-adjusted power in the α (median r = −0.18, p = 0.016), β (median r = - 0.04, p = 0.145) and γ (median r = 0.07, p = 0.773) bands (Figure S3). Therefore, we found that spontaneous and evoked EEG signals are associated with slow drift even after controlling for changes in 1/f noise [43–46].

## Discussion

In this study, we investigated if pre-stimulus α oscillations and the amplitude of the P1 component of the VEP could be utilized as an external signature of an internal brain state, a recently discovered neural activity pattern called slow drift [17]. We know from previous work that slow drift in macaque visual and prefrontal cortex is significantly associated with a host of eye metrics that are strongly indicative of changes in arousal [18]. Since pre-stimulus α power and P1 amplitude are also related to arousal, we wondered if a link could be established between a neural measure of an internal brain state acquired directly from the spiking activity of populations of neurons (i.e., slow drift) and indirect signals recorded non-invasively using EEG. Results showed that slow drift was significantly associated with a pattern that is indicative of changes in the subjects’ arousal levels over time. When pre-stimulus α power was low and P1 amplitude was high, pupil size, false alarm rate and saccade velocity increased, whereas microsaccade rate and reaction time decreased.

A variety of studies have shown that pre-stimulus α power and P1 amplitude are related to a behavioral pattern that is associated with arousal level. For example, decreases in the amplitude of pre-stimulus α oscillations are typically accompanied by improved performance on detection tasks [19–26] and decreased reaction time [35–38], whereas the opposite is true of P1 amplitude [19,25,39–41]. This motivated us to ask if pre-stimulus α power was negatively associated with P1 amplitude. In support of previous research [43], we found that decreases in pre-stimulus α power were accompanied by increases in P1 amplitude. This antagonistic effect might be related to functional inhibition. Evidence suggests that neuronal excitability is reduced when pre-stimulus α oscillations are relatively strong [48–53]. Since early VEP components are thought to be generated in an additive manner on top of spontaneous activity [54–57] inhibition of low-level sensory areas might lead to a reduction in P1 amplitude. Regardless of the underlying mechanism, a direct prediction of our results is that pre-stimulus α oscillations and P1 amplitude should be correlated in an opposite manner with a constellation of arousal-related variables.

In this study, we explored if pre-stimulus α oscillations and P1 amplitude were associated with various task performance and eye metrics that have been strongly linked to global changes in brain state. Results showed that pre-stimulus α power was negatively correlated with false alarm rate and saccade velocity, whereas it was positively correlated with reaction time. In contrast, P1 amplitude was negatively correlated with saccade velocity and microsaccade rate. Taken together, these findings suggest that the subjects’ arousal levels were changing slowly over time. As described above, decreases in pre-stimulus α power and increases in P1 amplitude are known to be associated with improved performance on detection tasks [19–26,39,40]. That these metrics were also coupled with increases in false alarm rate and saccade velocity as well as decreases in microsaccade rate and reaction time suggests that they can be taken as external measures of heightened arousal levels. In addition, our findings extend recent work in humans [31,32] showing that pre-stimulus α power is associated with changes in response criterion as opposed to sensitivity.

The principles of signal detection theory [27] dictate that improved performance on detection tasks can arise due to changes in sensitivity or response criterion [28–30]. Recent work has shown that decreased pre-stimulus α power is associated with increased hit rate and false alarm rate, which suggests that performance improvements linked to decreased pre-stimulus α power might occur due to changes in response criterion [31,32]. Our results support this hypothesis, as pre-stimulus α power was negatively associated with false alarm rate. Furthermore, we found that slow drift was positively correlated with hit rate (median r = 0.41, p < 0.001) and false alarm rate (median r = 0.53, p < 0.001) across sessions, consistent with previous research [17]. Luo and Maunsell (2015) found that changes in sensitivity, not response criterion were related to single-neuron and pairwise statistics in V4. In contrast, our results suggest that V4 activity is related to a shift in response criterion that was indexed by decreased pre-stimulus α power. One possible explanation for this discrepancy is that Luo and Maunsell (2015) manipulated response criterion in a spatially selective manner to avoid shifts arising due to global changes in arousal. In our study, response criterion was not manipulated directly, so shifts are likely to have occurred spontaneously due to global changes in the subjects’ arousal levels. In addition, we averaged across electrode sites because decreases in pre-stimulus α power/increases in P1 amplitude are associated with improved performance on detection tasks across large swathes of cortex [19,22,31]. These findings suggest that spatially selective and global shifts in response criterion might be governed by distinct neural mechanisms. This is an important issue to address, not just because it has the potential to resolve a discrepancy in the literature, but it also might explain why pre-stimulus α power on spatial attention tasks is associated with a pattern that is more akin to changes in sensitivity i.e. decreased/increased pre-stimulus α power over electrodes contralateral/ipsilateral to the attended hemifield [58].

In summary, we found that two commonly used metrics of cognitive state in human EEG studies, pre-stimulus α power and P1 amplitude, are associated with gradual shifts in the underlying population structure of neural activity throughout the brain. Together, these measures at the scalp, and in the cortex, were predictive of changes in the monkeys’ arousal levels over time. These findings show that indirect measures of neural activity can be used to index a global signature of arousal. They support recent work showing that pre-stimulus α power is associated with response criterion [59]. Finally, by linking a vast EEG literature in humans with simultaneous scalp/microelectrode array recordings in macaques our results bridge the gap between large-scale signals and the responses of single cortical neurons.

## Methods

### Subjects

Two adult rhesus macaque monkeys (*Macaca mulatta*) were used in this study. A previous report [60] presented analysis of different aspects of the same experiments described here. Surgical procedures to chronically implant a titanium head post (to immobilize the subjects’ heads during experiments) and microelectrode arrays were conducted in aseptic conditions under isoflurane anesthesia, as described in detail by Smith and Sommer (2013). Opiate analgesics were used to minimize pain and discomfort during the perioperative period. Neural activity was recorded using 100-channel “Utah” arrays (Blackrock Microsystems) in V4 (Monkey 1 = right hemisphere; Monkey 2 = left hemisphere). The arrays comprised a 10×10 grid of silicon microelectrodes (1 mm in length) spaced 400 μm apart. Experimental procedures were approved by the Institutional Animal Care and Use Committee of the University of Pittsburgh and were performed in accordance with the United States National Research Council’s Guide for the Care and Use of Laboratory Animals.

### Microelectrode array recordings

Signals from each microelectrode in the array were amplified and band-pass filtered (0.3– 7500 Hz) by a Grapevine system (Ripple). Waveform segments crossing a threshold (set as a multiple of the root mean square noise on each channel) were digitized (30KHz) and stored for offline analysis and sorting. First, waveforms were automatically sorted using a competitive mixture decomposition method [62]. They were then manually refined using custom time amplitude window discrimination software (code available at https://github.com/smithlabvision/spikesort), which takes into account metrics including (but not limited to) waveform shape and the distribution of interspike intervals [63]. A mixture of single and multiunit activity was recorded, but we refer here to all units as “neurons”. The mean number of V4 neurons across sessions was 70 (SD = 11) for Monkey 1 and 31 (SD = 16) for Monkey 2.

### EEG recordings

We recorded EEG from 8 Ag/AgCl electrodes (Grass Technologies) adhered to the scalp with electrically conductive paste. The electrodes were positioned roughly at the following locations: Fz, Iz, CP3, CP4, F5, F6, PO7, and PO8 (see Fig. 1B). Signals were referenced online to a steel screw on the titanium head post, digitized at 1 kHz and amplified by a Grapevine system (Ripple) and low-pass filtered online at 250 Hz. Segments of EEG data (a segment is defined as a 400ms stimulus period or a 300-500ms pre-stimulus period) were considered excessively noisy and removed if any of the channels had a standard deviation 10 times greater than the mean of the entire session. We also removed segments of data if any of the channels exhibited a flat signal defined as a standard deviation less than 300 nanovolts.

### Visual stimuli

Visual stimuli were generated using a combination of custom software written in MATLAB (The MathWorks) and Psychophysics Toolbox extensions [64,65]. They were displayed on a CRT monitor (resolution = 1024 × 768 pixels; refresh rate = 100Hz), which was viewed at a distance of 36 cm and gamma-corrected to linearize the relationship between input voltage and output luminance using a photometer and look-up-tables.

### Behavioral task

Subjects fixated a central point (diameter = 0.6°) on the monitor to initiate a trial (Figure 1A). Each trial comprised a sequence of stimulus periods (400ms) separated by fixation periods (duration drawn at random from a uniform distribution spanning 300-500ms). The 400ms stimulus periods comprised pairs of drifting full-contrast Gabor stimuli. One stimulus was presented in the aggregate receptive field (RF) of the recorded V4 neurons, whereas the other stimulus was presented in the mirror-symmetric location in the opposite hemifield. Although the spatial (Monkey 1 = 0.85cycles/°; Monkey 2 = 0.85cycles/°) and temporal frequencies (Monkey 1 = 8cycles/s; Monkey 2 = 7cycles/s) of the stimuli were not optimized for each individual V4 neuron they did evoke a strong response from the population. The orientation of the stimulus in the aggregate RF was chosen at random to be 45 or 135°, and the stimulus in the opposite hemifield was assigned the other orientation. There was a fixed probability (Monkey 1 = 30%; Monkey 2 = 40%) that one of the Gabors would change orientation by ±1, ±3, ±6, or ±15° on each stimulus presentation. The sequence continued until the subject) made a saccade to the changed stimulus within 400ms (“hit”); 2) made a saccade to an unchanged stimulus (“false alarm”); or 3) remained fixating for more than 400ms after a change occurred (“miss”). If the subject correctly detected an orientation change, they received a liquid reward. In contrast, a time-out occurred if the subject made a saccade to an unchanged stimulus delaying the beginning of the next trial by 1s. It is important to note that the effects of spatial attention were also investigated (although not analyzed in this study) by cueing blocks of trials such that the orientation change was 90% more likely to occur in one hemifield relative to the other hemifield.

### Eye tracking

Eye position and pupil diameter were recorded monocularly at a rate of 1000Hz using an infrared eye tracker (EyeLink 1000, SR Research).

### Microsaccade detection

Microsaccades were defined as eye movements that exceeded a velocity threshold of 6 times the standard deviation of the median velocity for at least 6ms [66–68]. They were required to be separated in time by at least 100ms. In addition, we removed microsaccades with an amplitude greater than 1° and a velocity greater than 100°/s. To assess the validity of our microsaccade detection method, the correlation (Pearson product-moment correlation coefficient) between the amplitude and peak velocity of detected microsaccades (i.e., the main sequence) was computed for each session. The mean correlation between these two metrics across sessions was 0.86 (SD = 0.07) indicating that our detection algorithm was robust [69].

### Pre-stimulus power

To calculate the mean amplitude of oscillations in different frequency bands during pre-stimulus periods the signals from all 8 EEG electrodes were averaged together. We adopted this approach because previous research has shown that pre-stimulus α power across a wide range of frontal, midline and occipital locations is associated with performance on detection tasks [19,22,31]. Furthermore, the overall pattern of results did not change when signals were averaged across frontal, midline and posterior electrodes. We then computed a FFT of the Hanning-windowed segment of data spanning the entire duration of the pre-stimulus period (300 - 500ms) and computed mean power in the α, β and γ bands. To isolate slow changes, the data for each trial and power band was binned using a 30-minute sliding window stepped every 6 minutes (Figure 2A).

### P1 amplitude

To calculate the VEP during stimulus periods the signals from all 8 EEG electrodes were averaged together. We adopted this approach for similar reasons to those described above i.e., because P1 amplitude across a wide range of frontal, midline and occipital locations is associated with performance on detection tasks [19]. In addition, the overall pattern of results remained the same when signals were averaged across frontal, midline and posterior electrodes. We then computed the VEP for the period spanning 200ms before to 200ms after stimulus onset, filtered the segment between 1 and 25Hz, and baseline-corrected by subtracting the average voltage for the period spanning 200ms before stimulus onset. To isolate slow changes over time, we divided the session into 30-minute bins stepped every 6 minutes and computed the mean VEP for each time bin. We were interested in whether slow drift was associated with the amplitude of the P1 component. Hence, we calculated the maximum amplitude of the peak occurring ∼100ms after stimulus onset from the VEP for each time bin (Figure 2B). Note that we were only able to reliably compute VEPs in 3 sessions for Monkey 2, which means the majority of evoked EEG data is from Monkey 1 (17 sessions).

### Aperiodic-adjusted pre-stimulus power

We wanted to rule out the possibility that slow drift was associated with gradual changes in 1/f noise [43–46]. Therefore, we carried out a supplementary analysis in which slow drift was correlated with aperiodic-adjusted power in different frequency bands. As before, signals from all 8 EEG electrodes were averaged together. We then computed a FFT of the Hanning-windowed segment of data spanning the first 300ms of the pre-stimulus period. To isolate slow changes over time, we divided the session into 30-minute bins stepped every 6 minutes and computed the mean FFT for each time bin. We then subtracted the aperiodic signal from the FFT for each time bin, which was computed by fitting an exponential function [46]. Finally, mean pre-stimulus power in the α, β and γ bands was calculated from the aperiodic-adjusted FFT for each time bin. Note that the frequency axis of the FFT for each pre-stimulus interval had to be identical in this analysis so as to reliably estimate the aperiodic signal. Since accounting for broadband changes in 1/f noise did not change the overall pattern of results (Figure S3) we decided to correlate slow drift with pre-stimulus power in different frequency bands using the full duration of the pre-stimulus interval (300-500ms) to enhance frequency resolution.

### Task performance metrics

Hit rate was defined as the number of saccades toward a stimulus that changed orientation divided by the total number of stimuli presented that changed orientation. False alarm rate was the number of saccades toward a stimulus that did not change orientation divided by the total number of stimuli presented that did not change orientation. To isolate slow changes, hit rate and false alarm rate were calculated in 30-minute bins stepped every 6 minutes. The width of the bins and step size were chosen based on previous research to ensure reliable estimates over a relatively small number of trials [17].

### Eye metrics

Mean pupil diameter was measured during stimulus periods, whereas microsaccade rate was measured during fixation periods between the visual stimulus presentations [18]. We did not include the initial fixation period when measuring microsaccade rate as there was an increase in eye position variability during this period resulting from fixation having been established a short time earlier (300-500ms). Such variability was not present in subsequent fixation periods. Reaction time and saccade velocity were measured on trials in which the subjects were rewarded for correctly detecting an orientation change. Reaction time was defined as the time from when the change occurred to the time at which the saccade exceeded a velocity threshold of 100°/s. Saccade velocity was the peak velocity of the saccade to the changed stimulus. To isolate slow changes in the eye metrics over time the data for each session was binned using a 30-minute sliding window stepped every 6 minutes.

### Calculating slow drift

The spiking responses of populations of neurons in V4 were measured during a 400ms period that began 50ms after stimulus presentation (Figure 4A). To control for the fact that some neurons had a preference for one orientation (45 or 135°) over the other residual spike counts were calculated. We subtracted the mean response for a given orientation across the entire session from individual responses to that orientation. To isolate slow changes in neural activity over time, residual spike counts for each V4 neuron were binned using a 30-minute sliding window stepped every 6 minutes (Figure 4A). PCA was then performed to reduce the high-dimensional residual data to a smaller number of latent variables [70]. Slow drift in V4 was estimated by projecting the binned residual spike counts for each neuron along the first principal component.

### Aligning slow drift across sessions

As described above, slow drift was calculated by projecting binned residual spike counts along the first principal component. The weights in a PCA can be positive or negative [47], which meant the sign of the correlation between slow drift and a given metric was arbitrary. Preserving the sign of the correlations was particularly important in this study because we were interested in whether slow drift was associated with a pattern that is indicative of changes in the subjects’ arousal levels over time i.e., decreased pre-stimulus α power and increased P1 amplitude. The method for aligning slow drift across sessions is described in detail by Johnston et al. [18]. Briefly, we projected evoked responses acquired during stimulus periods and spontaneous responses acquired during pre-stimulus periods onto the first principal component. The sign of the slow drift was flipped if the mean projection value for the evoked responses was less than that for the spontaneous responses i.e., if the relationship that was naturally observed for unprojected data did not hold true.

## Acknowledgements

M.A.S. was supported by NIH Grants R01 EY022928, R01 MH118929, R01 EB026953, and P30 EY008098; NSF Grant NCS 1734901; a career development grant and an unrestricted award from Research to Prevent Blindness; and the Eye and Ear Foundation of Pittsburgh. A.C.S. was supported by NIH grant K99 EY025768. R.S.S. was supported by NIH Grant R90 DA023426. The authors would like to thank Ms. Samantha Schmitt for assistance with surgery and data collection.

---

**Figure S1.**
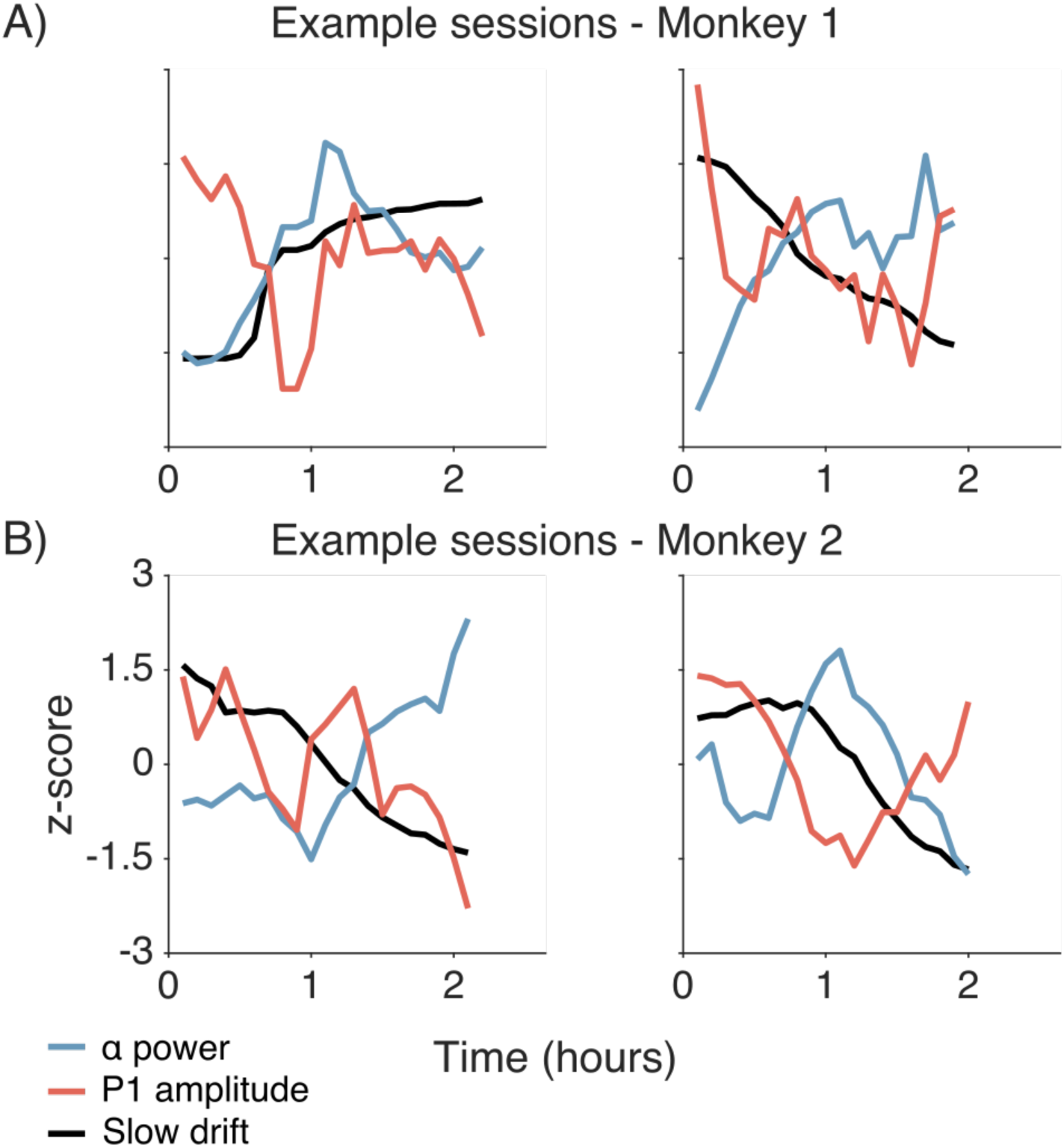
Relationships between slow drift, pre-stimulus α power and P1 amplitude (A) Example sessions from Monkey 1. Each metric has been z-scored for illustration purposes. (B) Same as (A) but for Monkey 2.

**Figure S2.**
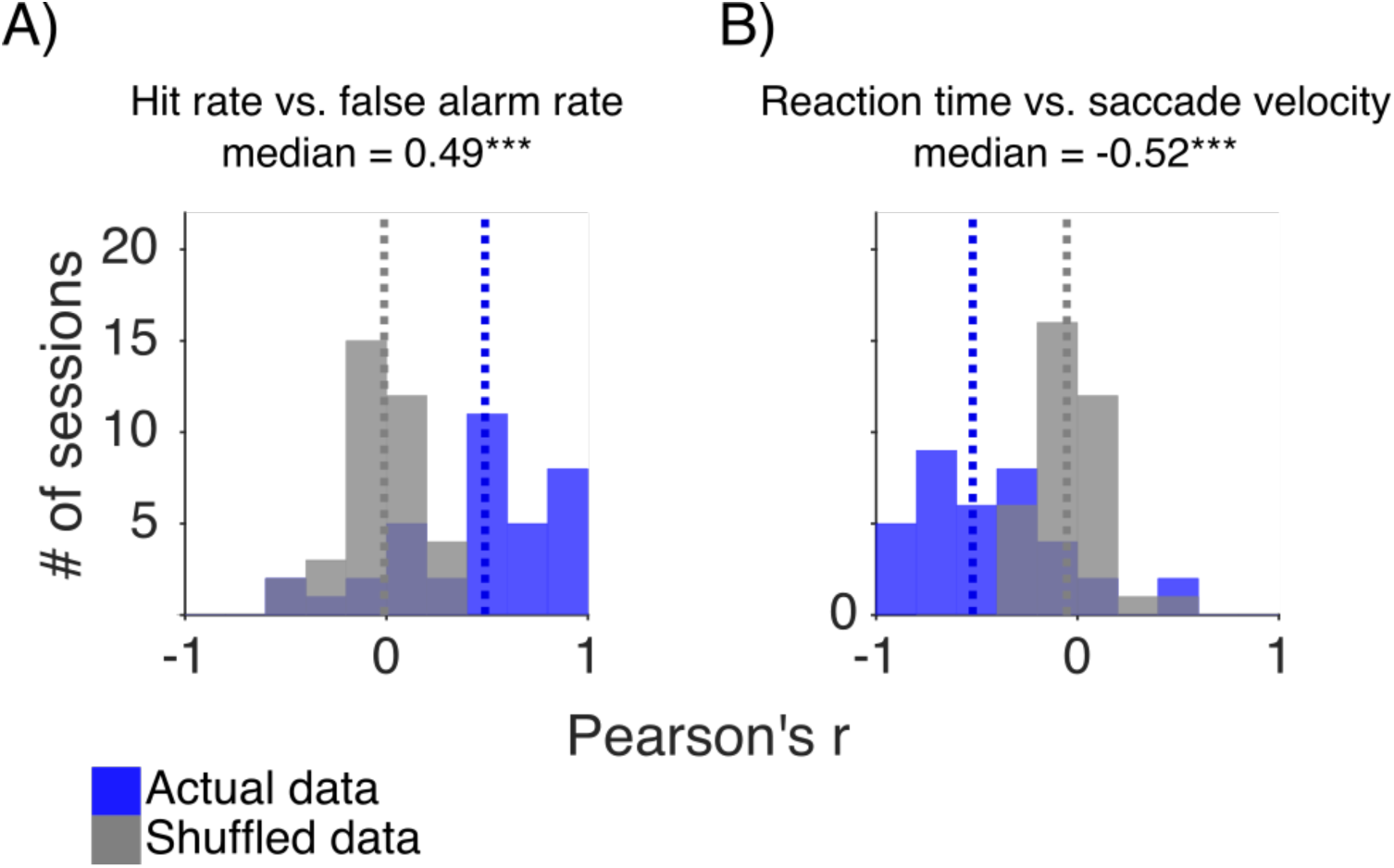
(A) Histogram showing distributions of r values (black) across sessions between hit rate and false alarm rate. (B) Same as (A) but for r values computed between reaction time and saccade velocity. In (A) and (B) actual distributions were compared to shuffled distributions (grey) using two-sided permutation tests (difference of medians). Median r values are indicated by dashed lines. p < 0.05*, p < 0.01**, p < 0.001***.

**Figure S3.**
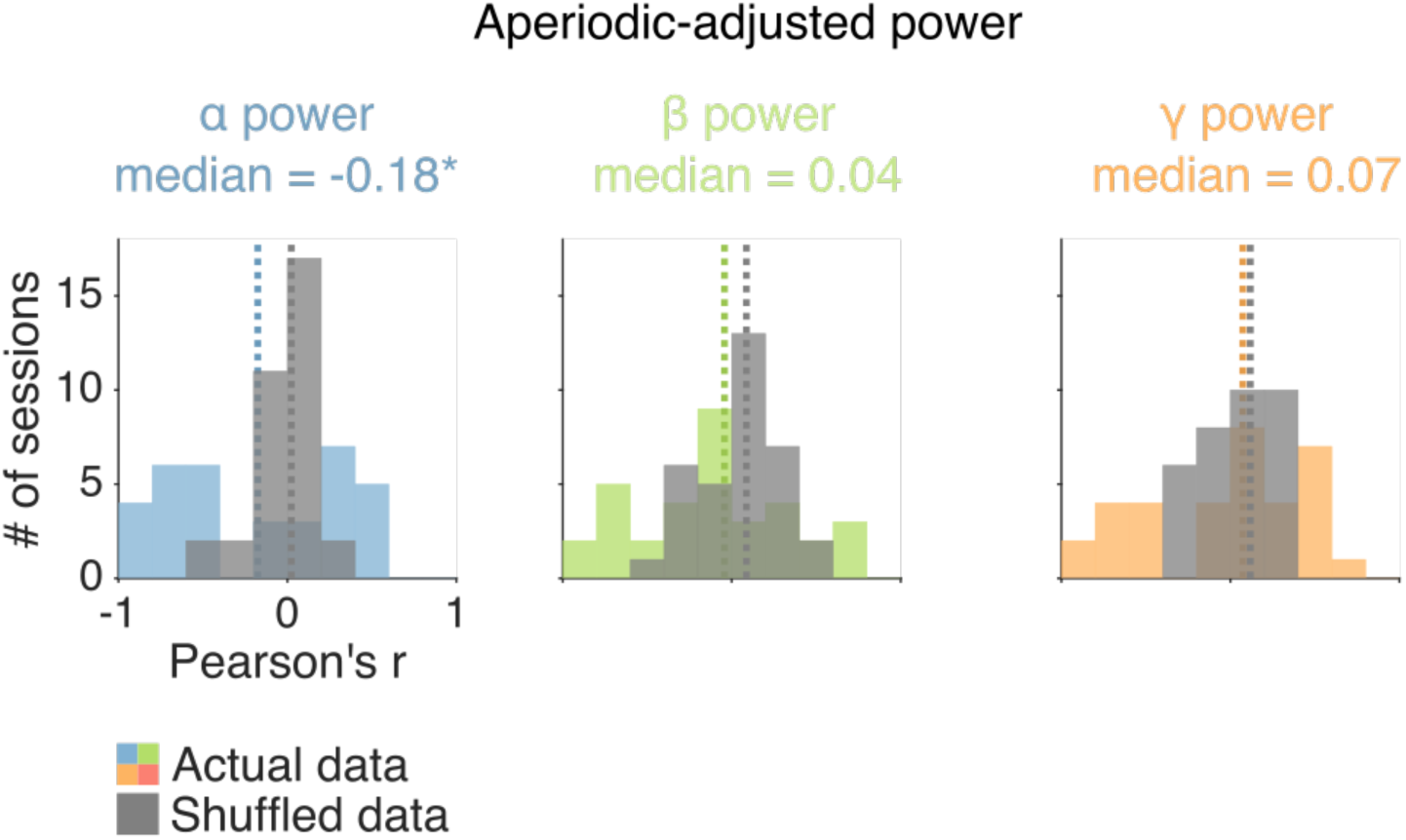
Correlations between aperiodic-adjusted power and slow drift. Histograms showing distributions of r values (colors) across sessions between slow drift and EEG power in different frequency bands. Actual distributions were compared to shuffled distributions (grey) using two-sided permutation tests (difference of medians). Median r values are indicated by dashed lines. p < 0.05*, p < 0.01**, p < 0.001***.

